# Crystal structure of Fatty Acid Thioesterase A bound by 129 fragments provides diverse development opportunities

**DOI:** 10.1101/2024.12.23.629660

**Authors:** Ekaterina Kot, Matteo P. Ferla, Patricia H. Hollinshead, Charlie W.E. Tomlinson, Daren Fearon, Jasmin C. Aschenbrenner, Lizbé Koekemoer, Max Winokan, Michael Fairhead, Xiaomin Ni, Katherine S. England, Laura Ortega Varga, Mark G. Montgomery, Nicholas P. Mulholland, Frank von Delft

## Abstract

To alleviate the growing issue of herbicide resistance, diversification of the herbicide portfolio is necessary. A promising yet underutilised mode of action (MoA) are Fatty Acid Thioesterases (Fat), which terminate *de novo* fatty acid (FA) biosynthesis by releasing fatty acids from acyl carrier protein (ACP) cofactors. These enzymes impact plant growth and sterility by determining the amount and length of FAs present. To find novel chemical matter targeting this MoA we have solved the crystal structure of *Arabidopsis thaliana* FatA to 1.5Å and conducted a crystallographic fragment screen which identified 141 hits. Our fragments recapitulated interactions made by current herbicides, but also targeted novel regions, namely the active site and the dimer interface. Ten fragments demonstrated on-scale potency, two of those exploiting different interactions to known herbicides. The screen also revealed large conformational changes exploited by some fragments, including stabilisation of a loop and shifts in the ACP-binding lid domain. These changes alter the accessibility of the substrate binding site by reducing the size of the apparent substrate access channel. Finally, the fragments readily lend themselves to designing compounds that both merge motifs and allow catalogue-based SAR exploration. Elaboration of one of the fragments resulted in an improvement of affinity from ∼20 μM to ∼90nM KD. Thus our data may facilitate accelerated early discovery of alternative inhibitors that are novel in both chemical scaffolds and modes of interaction.

## Introduction

Herbicide resistance is a growing issue threatening global food security. Out of the 31 biochemical processes targeted by herbicides, known as sites of action, 21 have had reported cases of resistance, equating to 168 different herbicides^1^. Herbicide resistance can be mitigated by sequential or combinatorial use of herbicides with different mechanisms of controlling the plants, known as modes of action. For that, herbicides must be developed that target novel or underutilised sites of action.

Fatty acid thioesterases (FATs) were first classified as modes of action in 2018 when BASF identified them as a target of cinmethylin, three decades after its release onto the market^2^. Since then, several more herbicides have been “posthumously” identified as targeting FATs and so in 2019, the herbicide resistance action committee (HRAC) publicised them as sites of action of cinmethylin and methiozolin; and in 2024, oxaziclomefone, bromobutide, cumyluron and tebutam were also added^3–5^. FATs offer an attractive MoA due to their absence from Animalia and a subsequently lower risk of toxicity. Moreover, FAT herbicides are being commonly used to control resistant grasses in pre-emergent applications^6^. As a result, in 2023, a new chemical motif, 1,8-naphthyridine, was introduced into play by Bayer^7^ which inspired several papers to employ scaffold hopping approaches to access novel inhibitors based either on 1,8-naphthyridine or existing herbicides such as methiozolin^7–11^. However, the pharmacophore profile of current herbicides is quite similar (Fig. 1(B)) and where a structure is available, they all bind in the same binding site and interact with a similar set of residues (Fig. 4(A-C))^5^. This increases the risk of resistance; reduced sensitivity towards cinmethylin has already been found in lines with non-target specific resistance to HRAC group 1 and 3 herbicides, albeit to levels below the field-rate dose^12^. The current herbicides can also suffer from a lack of crop selectivity, soil-dependent variability in efficacy and possess sub-optimal physical properties such as high volatility (cinmethylin) and low water solubility^13–17^. Additional chemotypes could present opportunities to improve on these qualities.

**Figure 1.**
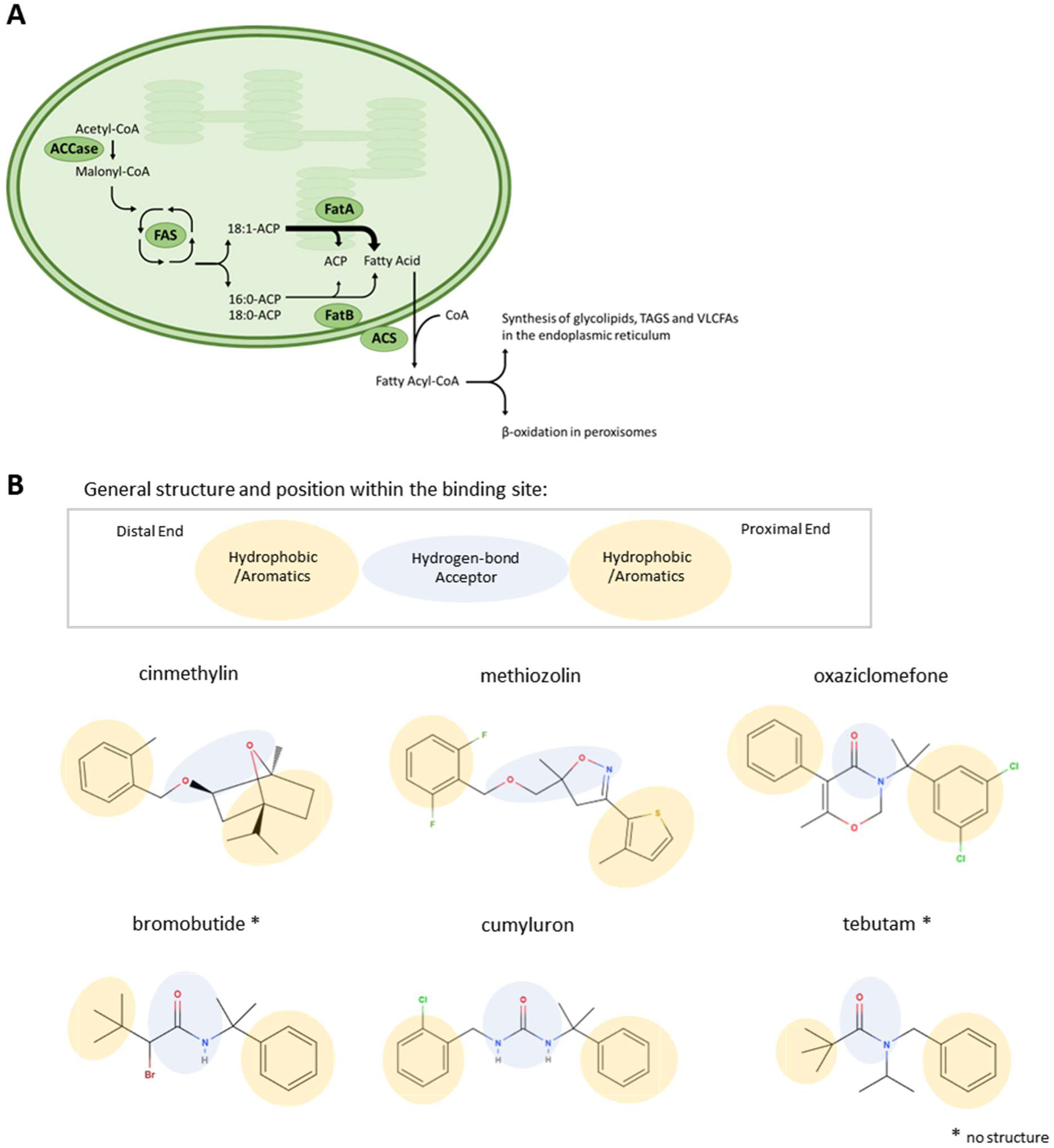
Biological pathway of FatA and its current inhibitors. **A.)** A simplified overview of fatty acid metabolism in the plastids of plants. CoA: Coenzyme A; ACCase: acetyl-Coenzyme A carboxylase; FAS: fatty acid synthase; Fat: fatty acid thioesterase; ACP: acyl carrier protein; ACS: acyl-CoA synthetase; TAGS: triacylglycerols; VLCFAs: very-long chain fatty acids. **B.)** Chemical structures of currently recognised herbicides targeting FatA highlighting similarity in their general pharmacophoric profile. Distal and Proximal Ends refer to orientation within the herbicide binding pocket based on their structures or estimates where no structure was available.

FATs are involved in the *de novo* biosynthesis of fatty acids (FAs) inside the plastids of higher plants by terminating the synthesis through cleavage of the thioester bond between a FA and an acyl carrier protein (ACP) cofactor (Fig. 1(A)). They play a key role in determining the quantity and type of FAs produced by plants, and in turn, affect plant growth and sterility^18,19^. There are two families of FATs: FatAs which have a preference for 18:1 FAs and FatBs which have broader specificity and a preference for saturated 18:0 and 16:0 chains^20,21^. The first published crystal structure of a plant thioesterase was FatB from *Umbellularia californica*, however recently three FatA structures from *Lemna aequinoctialis* bound to spirolactams have been released ^8,22^.

Since knockout of both enzymes is necessary for a lethal effect, it is thought that current herbicides must dually inhibit both FatA and FatB^2^. However, FatA has been more extensively tested owing to the simplicity of its purification and crystallisation, and its inhibition *in vitro* correlating well with *in vivo* outcomes^5,22^. We have therefore chosen FatA as a protein target for the development of novel chemical entities.

Crystallographic fragment screening can be a powerful way to discover novel starting points for future inhibitors. Because the size of the fragments is so small (usually <250Da), they explore the chemical space more efficiently and are more likely to be complementary to the target than the larger, more complex compound libraries used in traditional high-throughput screening^23,24^. Combined with the high-throughput screening facilities of XChem at Diamond Light Source (DLS), 1000s of fragments can be screened in a week to explore the molecular recognition capabilities of a protein target^25,26^. Several such campaigns have led to hit discoveries and hit-to-lead compound progressions^27–30^. Crystallographic screening also bears the advantage of supplying structural data which can be used alongside enzymatic and biophysical assays during rational drug discovery. It provides exact binding poses of the compounds, enables mapping of key interactions to specific residues and allows for comparison of conformational changes between different datasets, all of which can guide the follow-up compound design. The agrochemical field has been slow to adapt routine usage of fragment-based screening because of greater accessibility of *in vivo* phenotypic screening at the early stages of development when compared to pharmaceuticals. However, when trying to develop orthogonal opportunities, starting from smaller building blocks can be highly beneficial, hence our use of fragments in this campaign.

Here, we have applied crystallographic fragment screening to rationally explore novel chemical space against FatA to provide starting points for the discovery of an inhibitor with an alternative mode of action to the currently available herbicides. We have built on the in-house work from Syngenta to establish a robust crystal system suitable for an XChem campaign. We report a 1.5Å crystal structure of *Arabidopsis thaliana* FatA as well as a set of 129 fragment-bound structures, which provide 141 different starting points for future inhibitor development. 10 fragments already show on-scale potency in an enzymatic assay, with the top fragment having an IC50 < 15μM. Three binding sites of interest have been identified, each occupied by a diverse set of chemical scaffolds. Comparison of fragment-bound structures revealed that some ligands can induce large conformational changes within FatA and that the N-terminal half responsible for substrate binding displays much greater flexibility than the catalytic C-terminal half. As a result of this campaign, several diverse follow-up opportunities have emerged.

## Results

### The crystal structure of FatA reveals a dimer and adopts the same fold as FatB

A crystal structure of FatA from *Arabidopsis thaliana*, a widely used model organism, was obtained using the protocol previously established by Syngenta (Syngenta, personal communication). The construct includes a truncation at the N-terminus to remove the chloroplast transit peptide and a non-cleavable His-tag at the C-terminus. The previously established crystallisation condition consisted of MES and ammonium sulphate, which yielded consistent crystals that appeared after 1 day and diffracted around 1.7 Å. The structure was solved by molecular replacement using a crystal structure provided by Syngenta (Syngenta, personal communications).

Automatic data reduction pipelines at Diamond often attributed each crystal to both the I422 and the I222 space groups. Since the former results in a monomer within the asymmetric unit, we chose to process all of our data as I222. Processing the two protein chains separately allowed us to distinguish between the occasional conformational variability between the two chains, which is present in only some of the datasets. Moreover, we believe the crystallographic dimer we observe is in fact physiological as it maps faithfully onto the published FatB structure, which was shown to be physiological through mutations at the dimer interface^22^.

FatA from *Arabidopsis thaliana* adopted the same fold as FatB from *Umbellularia californica* (Fig. 2(B))^22^; the two possess a 42.4% sequence identity. The dimer contains two chains, each forming two tandem hot-dog folds. Each chain can be split into two halves which both form a hot-dog fold and are connected by a long linker region through the back of the beta-sheet (Fig. 2(A)). The N-terminal half forms the substrate-binding cavity while the C-terminal half contains the active site^22^. Residues within the previously proposed catalytic centre^22,31^ – Asp262, His266, Glu300, Asn264 and Asn269 – are located proximal to each other and in a way that does not contradict the possibility of their involvement.

**Figure 2.**
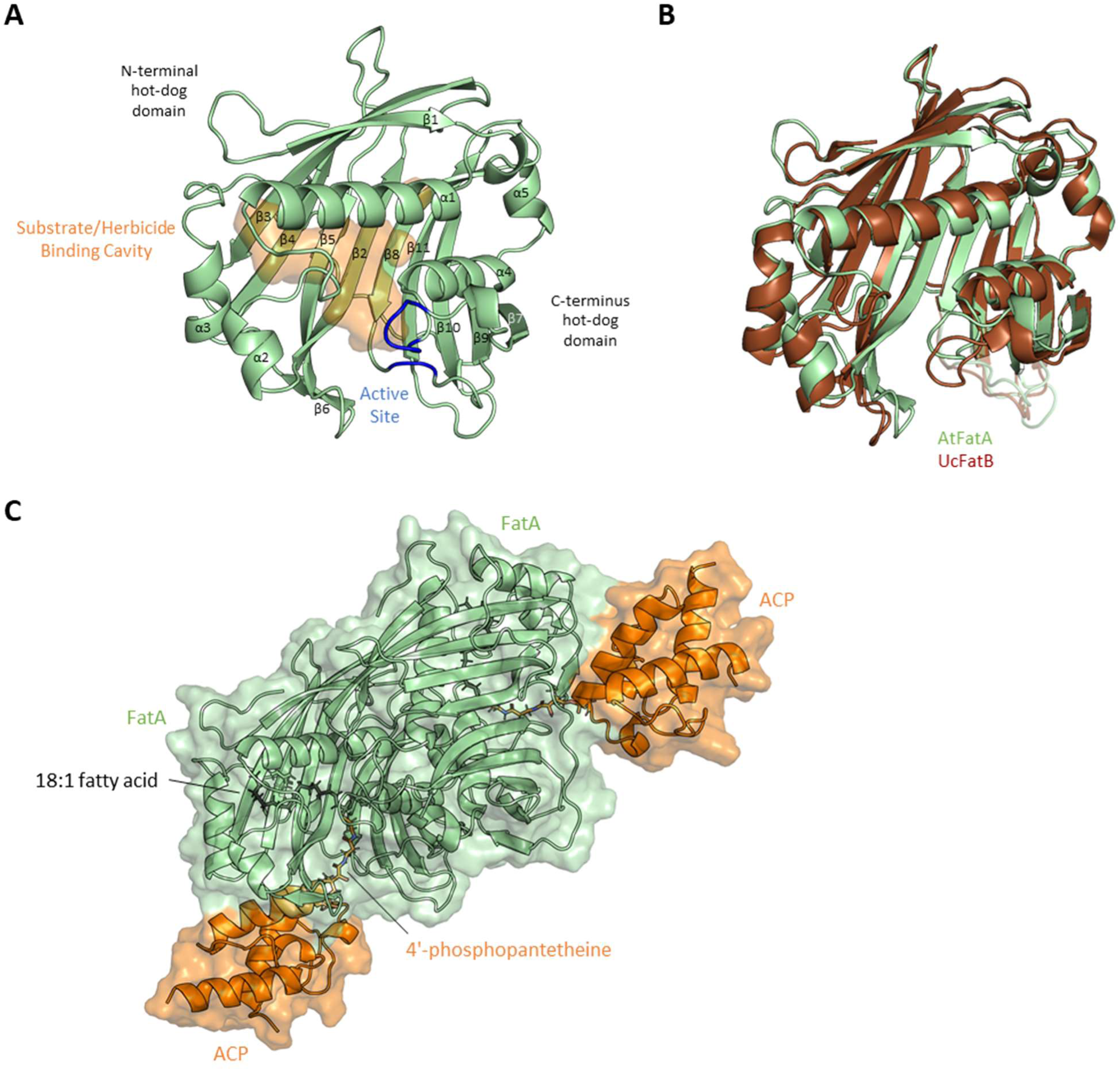
Structure of FatA and its biological assembly. **A.)** Crystal structure of Arabidopsis thaliana FatA (AtFatA). The substrate and herbicide binding cavity and active sites are shown in orange and blue respectively. α-helices and β-strands which make up two tandem hot-dog folds are labelled. **B.)** An overlay of FatA from Arabidopsis thaliana (AtFatA, PDB: 9HRQ) and Fata from Umbellularia californica (UcFatB, PDB: 5X04) crystal structures. AtFatA is shown in green; UcFatB is shown in brown. **C.)** A predicted model of ACP (shown in orange) bound to the crystallographic AtFatA dimer (shown in green) with an 18:1 fatty acid (shown in black) docked into the predicted substrate binding cavity.

Within the herbicide-binding site of FatA, we often observed a well-resolved density of an unidentified ligand (Fig. 5(F)). We postulate this may be a metabolite from the *E.coli* expression host since it does not resemble any component of the protein buffer or the crystallisation condition. However, upon soaking with DMSO as part of the solvent test, the density disappeared, thus we went ahead with a crystallographic fragment screen. An example of a structure containing the unidentified ligand, x0045, has been deposited in the PDB under ID 9HRQ, while the *apo* structure of FatA was deposited under ID 9HRR.

### Model of FatA’s biological assembly informs regions of importance and the mechanism of inhibition of known herbicides

The FatA-FA-ACP complex was modelled using our FatA crystal structure, ColabFold and a related *E.coli* AcpP structure crosslinked to a substrate mimic (Fig. 2(C))^32^. Based on this model, the FatA-ACP interface is formed of two loops between β3-4 and β8-9, a strip of the β-sheet perpendicular to the strands, a portion of the long linker between N- and C-terminal halves and residues 205-209 which connect to the lid domain formed of residues 197-206. The substrate was modelled as a possible transition state where it is covalently bound to Asp262. The proximity of His266 supports its role in activating a water molecule during hydrolysis. The long fatty acid fits well into the internal hydrophobic cavity of FatA, which is largely occluded from the outside with the access point between the lid domain and the active site. The double bond within the FA lines up to be within 4 Å of NE of Arg176, whose presence was perplexing within such a hydrophobic cavity. However, since this arginine is also present in FatB it is unlikely to contribute to the specificity of FatA towards 18:1 FAs.

### 129 fragments were found to bind FatA during a crystallographic fragment screen

A crystallographic fragment screen was conducted at Diamond’s XChem facility. Two crystallisation conditions were tested in a small 100-fragment pre-screen: the one used to obtain the *apo* structure (consisting of MES and ammonium sulphate) as well as one using cacodylate as a buffer instead. No improvement in hit rate and crystal survivability was observed with cacodylate, so the main screen used MES as the crystallisation buffer, however, hits unique to each condition were observed (see Supplementary Data).

For the main screen, two libraries (Enamine Essential Fragments and DSi-Poised^33^) were used yielding a total of 1101 compounds. The PanDDA algorithm identified binding events in 242 datasets. 129 structures contained fragments bound to pre-determined sites of interest and thus were processed further, giving rise to 141 individual hits (Fig. 3(A)). The hit number is greater since the fragments which bound simultaneously at different sites within the same crystal constitute different starting points for further development and thus were counted separately. Fragments bound outside the sites of interest were termed “misses” and not processed further.

**Figure 3.**
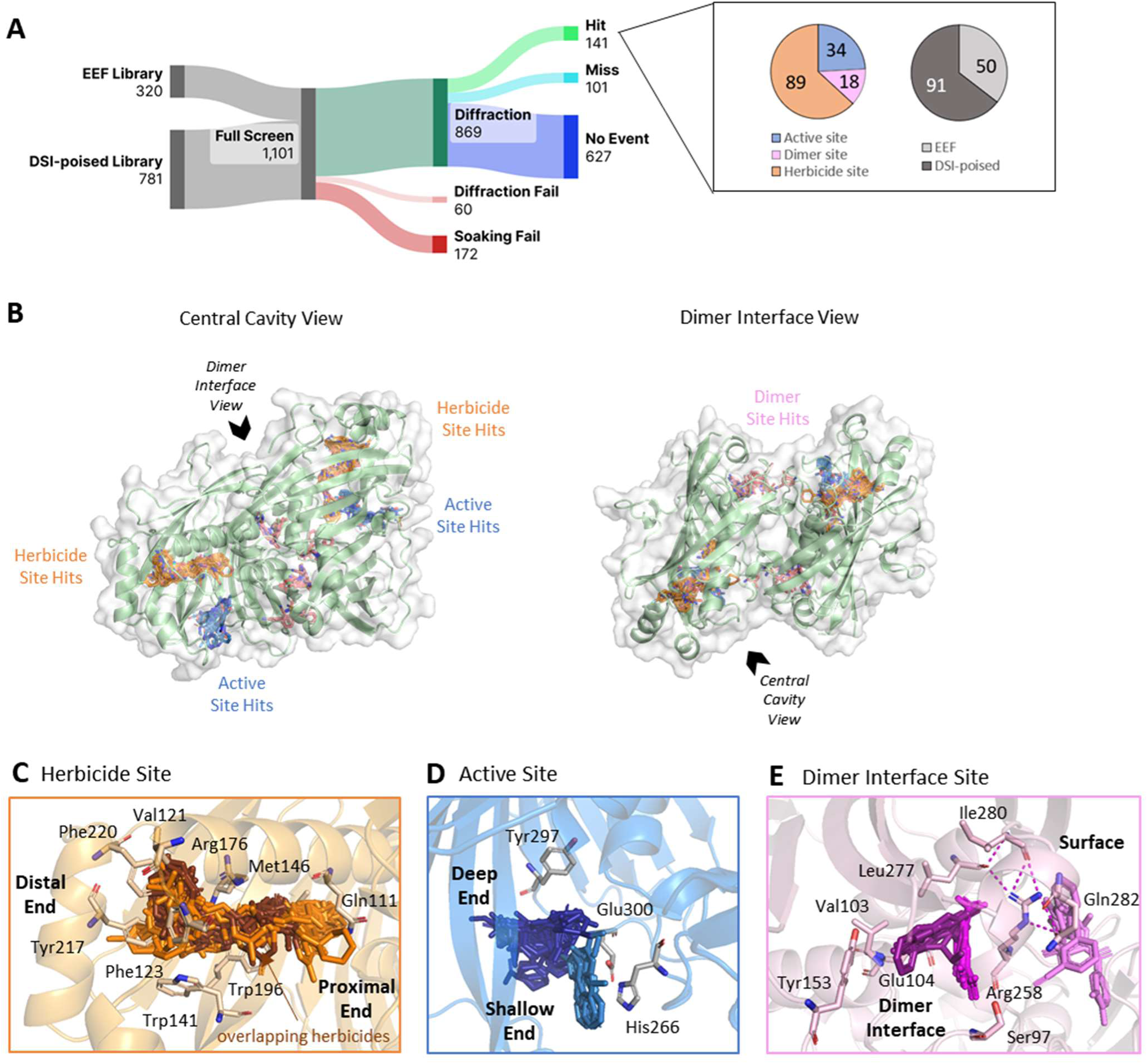
Overview of the crystallographic fragment screen on FatA and the distribution of hits across the three binding sites of interest. **A.)** A Sankey diagram of the crystallographic fragment screen of FatA with pie charts showing which libraries the fragments originated from and the distribution of hits across the sites of interest. EEF: Enamine Essential Fragment; DSI-poised: Diamond-SGC-iNEXT poised. Sankey diagram produced by SankeyArt.com. **B.)** Structure of the FatA dimer bound by all fragment hits. Close-ups of the **C.)** herbicide **D.)** active and **E.)** dimer interface sites showing frequently interacting residues. Where a site has been split into groups, these groups are labelled. For the herbicide binding site, atoms which are within 1Å of cinmethylin, methiozolin or oxaziclomefoneare coloured brown.

There was a 23% attrition rate composed of soaking fails where the crystal could not be mounted after soaking because of its deterioration and diffraction fails where the successfully mounted crystals did not diffract.

The three sites of interest were the herbicide binding site – based on the previously determined structures of commercial herbicides shown to target FatA, the active site – based on the position of residues previously shown to be important for catalysis, and the dimer site – located at the interface between the two protein chains chosen because formation of a dimer has been shown to be necessary for catalytic activity of plant FATs (Fig. 3(B))^5,22,31^. However, since targeting the dimer interface may result in stabilisers of the enzyme, this site carries the highest risk.

All fragment hit bound structures were deposited in the PDB under the group deposition ID G_1002328.

#### Fragments bound at the same site as commercial herbicides recapitulate key interactions as well as explore novel areas of the cavity

We name the herbicide binding site based on the fact that known commercial herbicides cinmethylin, methiozolin, and oxazioclomefone bind here in a similar way (Fig. 4(A-C)) (herbicide-bound structures provided by Syngenta, private communication). Moreover, this is the same site that our ACP-bound model suggests the natural substrate occupies during catalysis, making it likely that inhibitors bound here act by sterically excluding the substrate. Since the cavity is quite large, the side closest to the active-site was termed “proximal” while the end of the binding pocket was termed “distal” (Fig. 3(C)).

**Figure 4.**
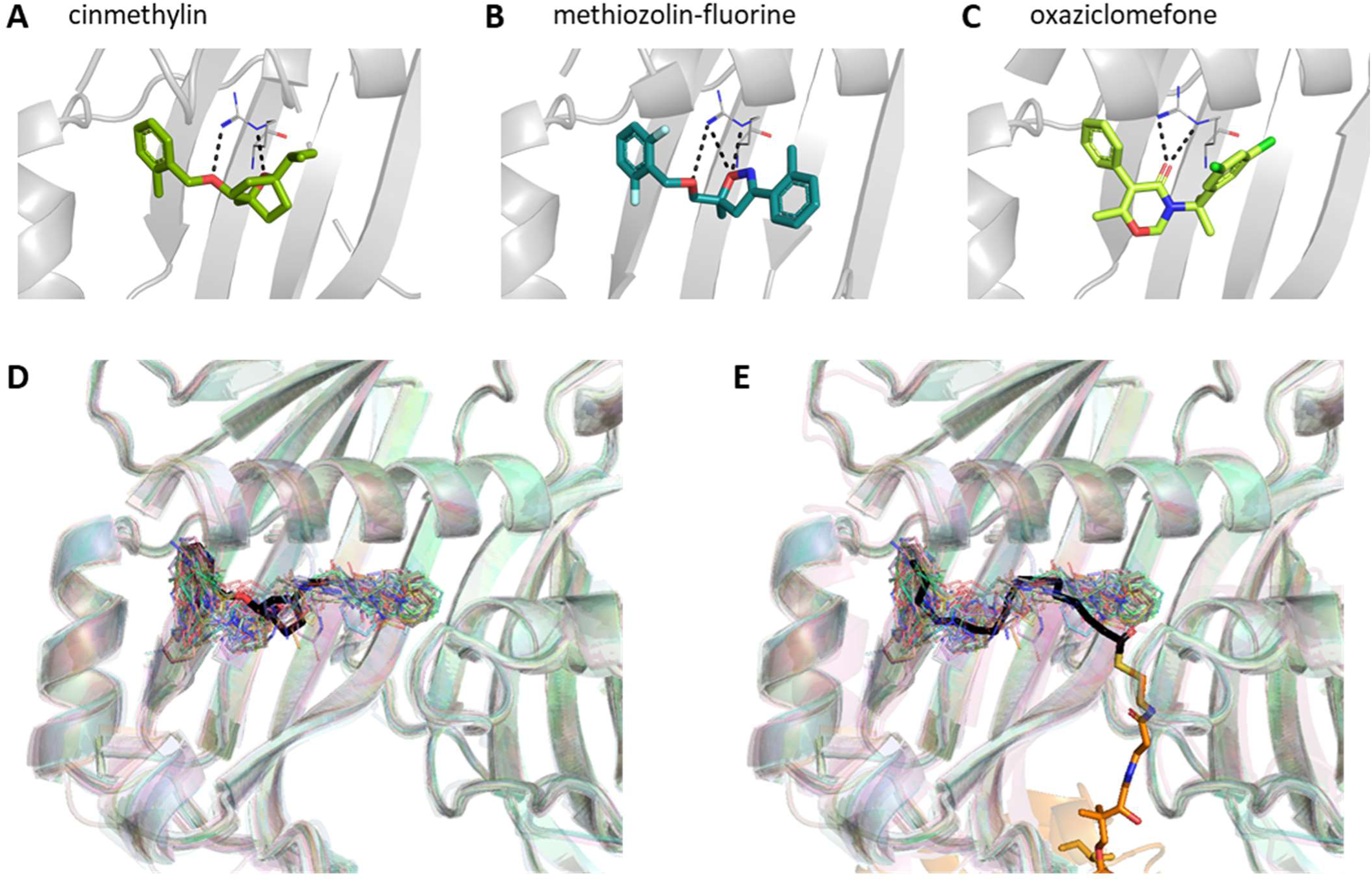
Fragments bound at the herbicide site overlap with commercial herbicides and the natural substrate. Crystal structures showing bound herbicides **A.)** cinmethylin (PDB: 9GRR), **B.)** methiozolin analogue methiozolin-fluorine (PDB: 9HMT) and **C.)** oxaziclomefone (PDB: 9GS1). An overlay of all hit-bound structures of FatA showing all fragments which bound at the nerbicide binding site. An alignment with **D.)** a crystal structure of cinmethylin and **E.)** modelled structure of 18:1-ACP bound FatA are shown.

The key defining residue of this herbicide site is Arg176 which hydrogen bonded to all the known FATs inhibitors and also aligns with the double bond of the 18:1 FA substrate in our model. FATs herbicides also form pi-pi and hydrophobic interactions with Val121, Phe123, Tyr217, Phe220, Trp141 and Trp196 via their hydrophobic motifs on either side of the H-bond acceptor.

We therefore compared the binding poses of our 89 fragments to that of the herbicides and the natural substrate (Fig. 4(D-E)). All the interactions made by current herbicides were recapitulated by subsets of our fragments. Tyr217 and Phe123 were especially frequent sites of interaction with aromatic motifs frequently observed overlaying the aromatic portion of the herbicides (Sup. Fig. S1). However, our fragments explored a larger proportion of the substrate binding cavity, especially the proximal half. Gln111 and Met146 located here were especially popular hydrophobic interactors. Gln111 also frequently formed H-bonds via its side-chain such as with x1285.

#### Some fragments bound at the active site make direct interactions with the catalytic histidine

The active binding site is composed of four loops. Two loops formed of residues 266-269 and 298-300 are part of the C-terminal half and contain the residues shown to be involved in catalysis. The other two loops are from the N-terminal half: the tip of the 122-132 loop and residues 197-206 which form the lid domain. Val142 which is located on the end of a beta-strand nearby also forms some interactions with the fragments.

The fragments bound at the active site were split into two groups: the “deep” half bound between β2 and β8 strands and the “shallow” half bound closer to the active site loops (Fig. 3(D)). Fragments in the deep half commonly interacted with Tyr297, which, while noncatalytic, is still proximal to the active site (Sup. Fig. S1). It formed both hydrophobic interactions and H-bonds via its backbone oxygen, for example, to x1891.

Within the shallow half of the catalytic residues, His266 and Asn269 make the most frequent interactions with the fragments, while Asn264 and Asp262 were largely uninvolved (Sup. Fig. S1). Since Glu300 has weak density, the reliability of interaction analysis is reduced. His266 was observed interacting via π-stacking with aromatic motifs of 5 unique fragments, all of which contain two heterocyclic rings, fused in most (4/5) cases (Sup. Fig. S2).

#### Fragments bound at the dimer interface cluster around an arginine involved in dimer interactions

The dimer site consists of a few fragments bound on the surface and a larger number within the dimer interface near Arg258 (Fig. 3(B,E)), which has previously been shown to be key in maintaining the dimer interface via a hydrogen bond network^22^. In the case of FatA, Arg258 forms H-bonds with backbone oxygens of Leu277 and Ile280, and side-chain of Gln282. Several of our fragments interact directly with this network: x1231 and x1236 bind Arg258 via π-cation interaction and an H-bond, respectively. Peripherally bound x1174 makes an Ar-HBond to Gln282 (Fig. 3(E)).

Those that bind within the dimer interface pocket do so in two groups which are at a 60° to each other but overlap on one end and can be distinguished by their pattern of interactions (Fig. 3(E)). Those closer to the surface can form H-bonds with Ser97 such as x1211, while the more buried group commonly interact with Val103 either hydrophobically via its side chain or form an H-bond with its backbone NH. Interactions with Tyr153 are also exclusive to this group and highly versatile; we observed H-bonding, π-stacking and hydrophobic interactions depending on the fragment bound. At the point where both groups overlap, both sets of fragments can form H-bonds with the side chain of Glu104.

### Binding of some ligands induces large, asymmetric movement

Upon binding of some ligands, large portions of the N-terminal hotdog domain of one chain, chain B, move closer to the C-terminal half relative to the *apo* state. Specifically, two regions: residues 122-142 which consist of a loop, helix 2, loop and beginning of β2, and residues 195-220 which begin as β5, form a large linker which sometimes contains β6 and finishes as a helix 3 (Fig. 5(A)) move 2.3Å and 3.3Å respectively. Both regions are proximal in space despite being part of different regions of the sequence, wrapping around the end of the substrate/herbicide binding site within the N-terminal half of the chain. Concurrently, these areas displayed higher B-factors compared to the rest of the structure, suggesting they remained somewhat dynamic after movement (Sup. Fig. S3).

**Figure 5.**
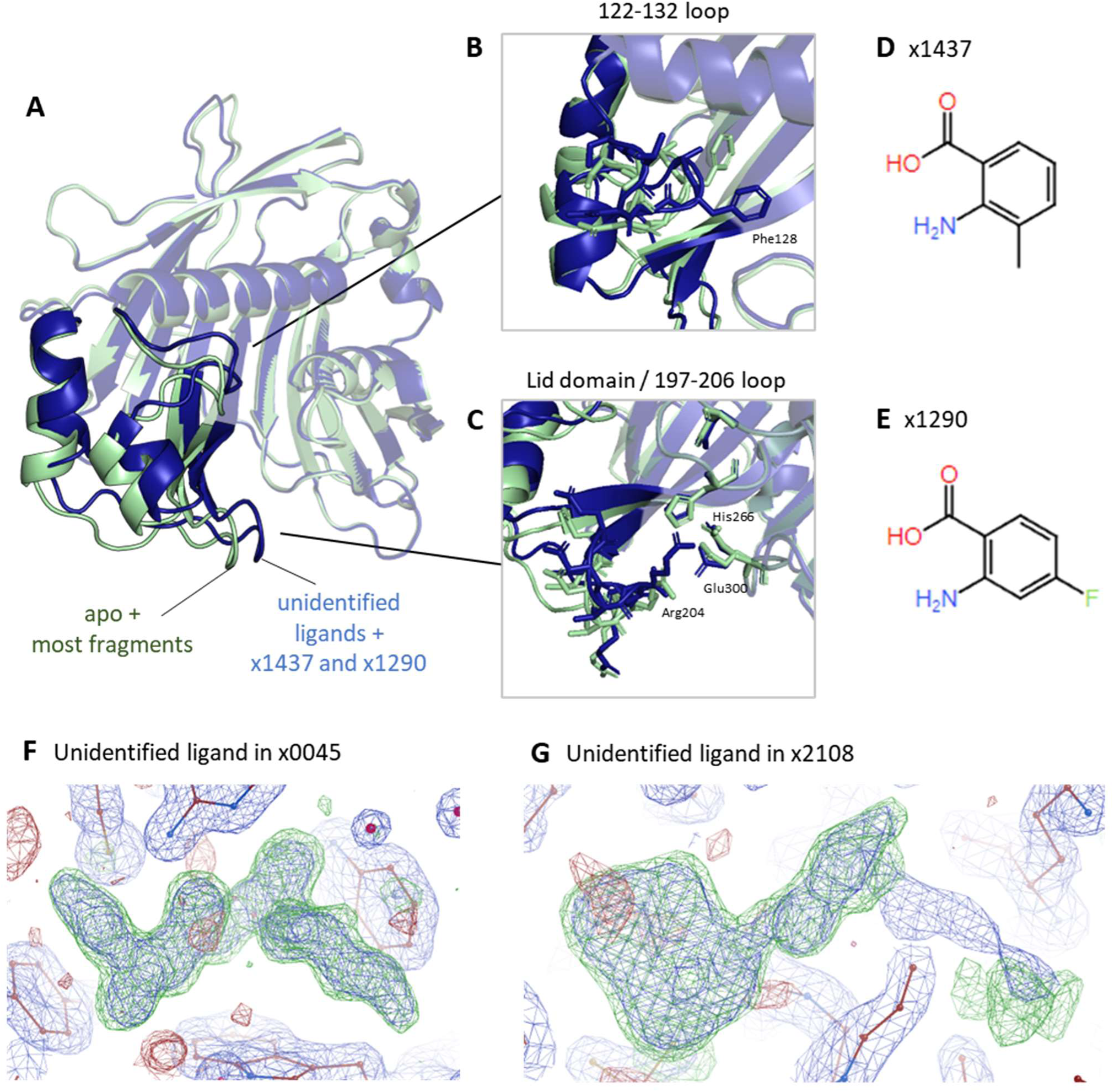
Large conformational changes within the N-terminal half of FatA occur upon binding of some ligand. **A.)** Overview of conformational changes induced from an overlay of x1163 and x143 7 crystal structures. Two regions of interest are identified-122-132 loop and a lid domain. **B.)** A close up of 122-132 loop movements especially the position of Phe128. C.)A close up of the lid domain/197-206 loop movement towards the active site loop. These changes were induced by fragments D.) x1437 and E.) x1290, F.) unidentified density in crystal x0045 obtained during crystallisation prior to the fragment screen, G.) unidentified density in chain B of x2108.

Specifically, we found four chemical entities which caused these significant shifts within chain B – all bound at the herbicide binding site: first, the unidentified ligand which was discovered during attempts to solve the *apo* structure, x0045, (Fig. 5(F)); secondly, within the dataset x2108, the herbicide binding site of chain B is occupied by another unidentified density which doesn’t resemble the expected fragment (captured at the chain A) (Fig. 5(G)); finally, two fragments from datasets x1290 and x1437 which share a similar chemical motif and are both bound at the herbicide binding site both caused the same shift (Fig. 5(D-E)).

Another dataset, x2025, was captured in an intermediate position between the majority of fragments and the four discussed above (Sup. Fig. S4). It doesn’t resemble fragments from x1290 and x1437 and its loop 122-132 is in a different conformation, thus it is closer to the rest of the fragments than the outliers.

Two regions of functional importance were identified to be contained within the mobile region. The 122-132 loop, which might form a gate to the substrate binding cavity via a bulky Phe128, and the 197-206 loop, which has previously been identified as a lid region, mediating interactions with ACP (Fig. 5(B-C)). All the ligands which caused the movements also led to the 122-132 loop becoming structured in chain B (where the ligand density is the strongest) and position Phe128 in an identical conformation which we defined as conformation A (Fig. 6(B)). The large conformational movements also caused the tip of the 122-132 loop and the lid domain to move closer to the active site loops which reduces the hole which leads to the substrate binding site and through which the substrate would have to go in order to align its thioester bond for catalysis^22^.

**Figure 6.**
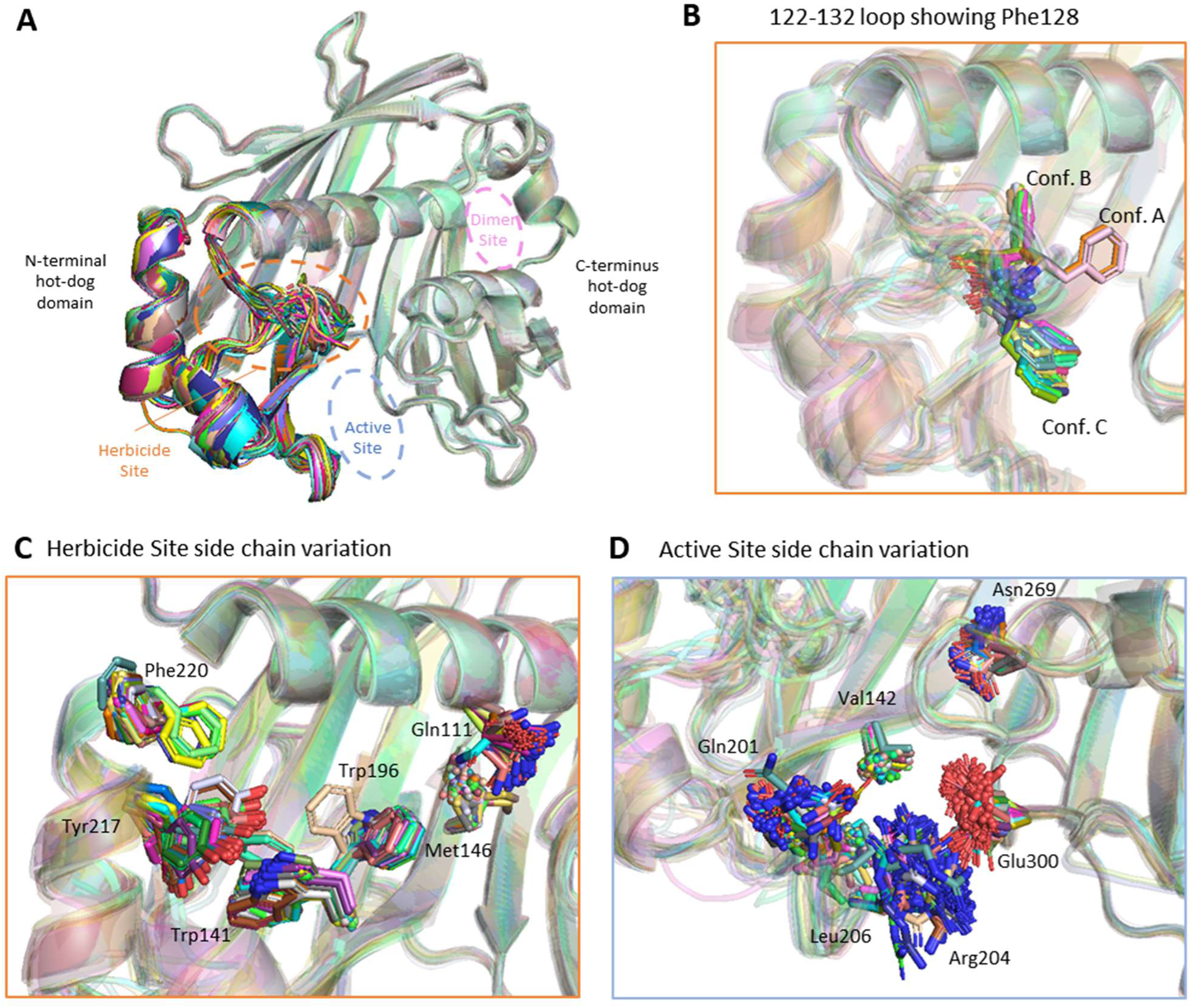
N-terminal half of FatA exhibits higher conformational flexibility than the C-terminal half as seen through the position of the 122-132 loop and side chains of the herbicide and active binding sites. **A.)** Overview of conformational flexibility from an overlay of all fragment bound structures. The herbicide and active sites located within the dynamic regions are highlighted in orange and blue respectively. **B.)** A close up of 122-132 loop flexibility as seen through three possible positions of Phe128. Side chain movements inside the **C.)** herbicide and **D.)** active sites. Specific chains which show particularly large conformational flexibility are shown and labelled.

### Substrate-binding N-terminal half of FatA displays much greater conformation variability than the catalytic C-terminal half

When comparing the rest of the structures obtained during the fragment screen which did not cause the large conformational movements outlined above, we noticed that the N-terminal hotdog domain of FatA, which is responsible for substrate binding and specificity, displays a greater degree of conformational variability when compared to the rest of the protein (Fig. 6(A))^20^. This affected the herbicide binding site and the active site, while the dimer site, despite being formed of residues from both A & B chains of FatA, showed little crystal-to-crystal variability.

#### A loop adjacent to the herbicide binding site displays a great deal of conformational variation

The 122-132 loop which forms a part of the substrate binding site was observed in a multitude of different conformational states (Fig. 6(B)). Specifically, residues 124-129 were often too disordered for the density to be modelled. However, upon binding of certain fragments, these residues became sufficiently ordered to be modelled. Comparison of the modelled loops revealed three main conformations which we defined by the position of Phe128 designated conformation A, B, and C. An analysis of the RMSD between the crystal structures corroborated these observations (Sup. Fig. S3). As mentioned above, conformation A was only observed together with large conformational changes. Another conformation, B, is predicted to clash with a modelled 18:1 FA and thus would cut off access to the substrate-binding cavity, offering an additional mechanism of inhibition for the fragments and herbicides.

Side chains of residues which lie on this loop, such as Phe123, Ala129 and Thr130, were captured in a variety of conformations. The low density and subsequently high B-factors reflect the relative mobility of these residues and the rest of the loop. Of those, Phe123 frequently interacts with fragments bound at the herbicide site despite the side chain being observed in a variety of conformations.

#### Residues involved in key interactions within the herbicide binding site displayed high conformational variability

Many of the residues located at the herbicide binding site adopted several conformations (Fig. 6(C)). Tyr217, which makes the most interactions with fragments out of all residues and binds some herbicides, was observed in several conformations. However, the density was well resolved in each case, suggesting that, unlike the residues of the 122-132 loop, Tyr217 adopts stable poses. Similarly, Phe220 was observed in two different conformations, again with good density to support each. Both of these residues are located on a α-helix 3, which moves during the global movement.

Two tryptophan residues which line the substrate binding cavity, Trp141 and Trp196, were generally observed in the same position, but in rare instances their side chains were flipped 90 degrees (Fig. 6(C)). Specifically, the fragments x1620 and x1747 which both contain a phenylmethoxy group stabilise the flipping of the Trp141 which brings the end of its ring within 4Å of the fragments. x1620 also stabilised a flipped conformation of Trp196.

Some of the variability resulted from poorly resolved density of side chains such as in the case of Gln111, Met146 and Thr174. The flexibility of these side chains may promote their interactions with a greater set of fragments. Two cysteines, Cys115 and Cys166 were observed and modelled in two side chain conformations.

#### The active binding site possesses lots of flexible side chains

High side chain flexibility within the active site was observed, both from residues on the N-terminal loops, such as Arg204, Leu201, Leu206 and Val142 and from the C-terminal half. Of the catalytic residues, Glu300 and Asn269 displayed particularly reduced densities indicative of movement (Fig. 6(D)). Because of this ambiguity, they have been modelled in a variety of conformations, each with reduced certainty. Such conformational flexibility may be necessary for the accommodation of the large diversity of chemistry we observed to bind here; some bound fragments would clash with side chain conformations observed in other datasets.

### Measurable inhibitory activity is already displayed by some fragments

64 fragments were available for testing in an *in vitro* enzyme assay. In this assay, the ACP cofactor was replaced by CoA and a thiol-reactive fluorescent dye was used to quantify the enzyme’s activity. 10 fragments were found to have on-scale potency with the top compound achieving a 13.0 μM IC50 (Fig. 7, Sup. Fig. S5). One of these fragments, x1214, produced a PanDDA event but did not result in convincing fragment density during refinement, leaving its binding site ambiguous. One fragment, x1944, did not demonstrate detectable inhibition, but an analogue where the pyrrolidine ring was substituted with an azepane to produce x1944*, had an IC50 of 43.9μM (Fig. 7(A)). Most of these fragments bound at the herbicide binding site and produced clear electron density.

**Figure 7.**
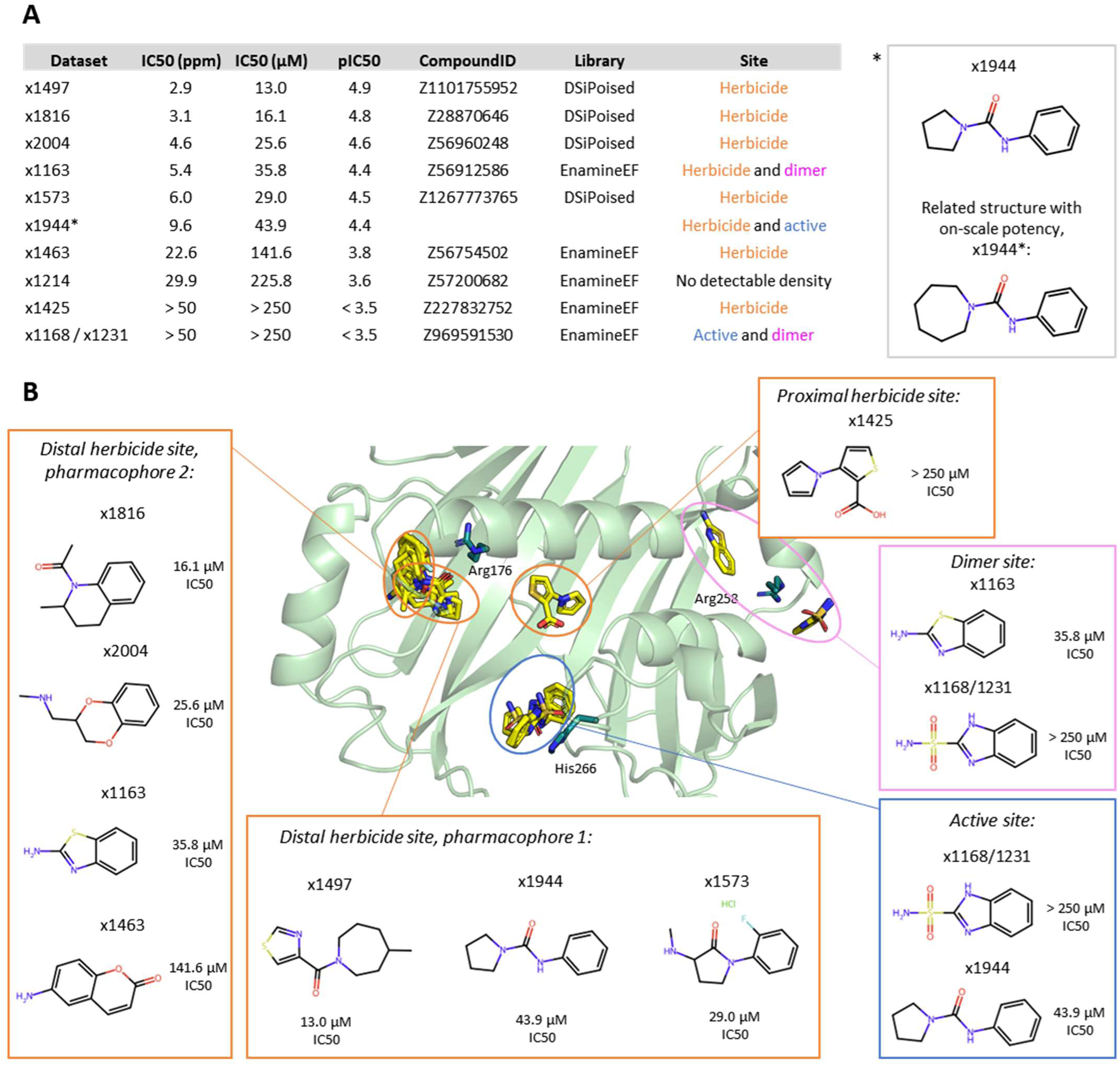
10 fragments have on-scale potency when tested in an enzymatic assay with FatA. **A.)** A summary table of the identity and inhibitory activity of the10 fragments. **B.)** The binding poses of the 9 fragments and their chemical structures.

x1168/x1231 bound at the active site and produced detectable activity, however, it was not possible to extract an exact IC50 value. It formed a π-stacking interaction with catalytic His266 and a water bridge with Asn269.

Of those bound at the herbicide binding site, two main pharmacophores were observed (Fig. 7(B)). Pharmacophore 1, consisting of x1497, x1573 and x1944, resembled the current herbicides; they formed a hydrogen bond with Arg176 while also possessing hydrophobic entities on either side of the hydrogen bond acceptor to interact with the greasy cavity. Pharmacophore 2, consisting of x1816, x2004, x1163 and x1463, resembled 1,8-naphthyridine: these were bicyclic, aromatic compounds which bound closer to the deep end of the pocket. While no interaction was common to all four, Val121, Phe123, Arg176 and Phe220 made interactions with three of the compounds. These were mainly hydrophobic, with occasional π-stacking and π-cation interactions via the rings. The last fragment, x1425, bound within the proximal end of the cavity and made both hydrophobic and H-bonding interactions with Gln111 and Asn269.

### The fragment screen offers several diverse opportunities for elaboration

Because of the disjointed nature of the three binding sites, separate elaboration of each is required. Fragment growing, merging and linking are the common strategies for fragment progression. As far as elaboration hypotheses, three principal ways of inquiry have emerged: elaboration based on conformational changes, prioritisation of compounds with measurable inhibition activity and/or maximisation of the number of key interactions.

A limited elaboration campaign based on growing of the active x1816 fragment was conducted and improvement in binding affinity was observed by SPR. Close analogues of x1816 were used to explore the binding capabilities of the herbicide pocket as well as tolerance to heterocyclic ring size. 60 were chosen based on docking results and either purchased from Enamine REAL or synthesised in-house. Of those, 27 had improved affinity when compared to the parent fragment which had a 17-20μM KD. The top compound, x1816-FU1 had a ∼90nM KD constituting a x220 improvement from a single design-make-test-analyse cycle. Structures and affinities of the top 10 follow-up compounds are shown in Figure 8(A).

**Figure 8.**
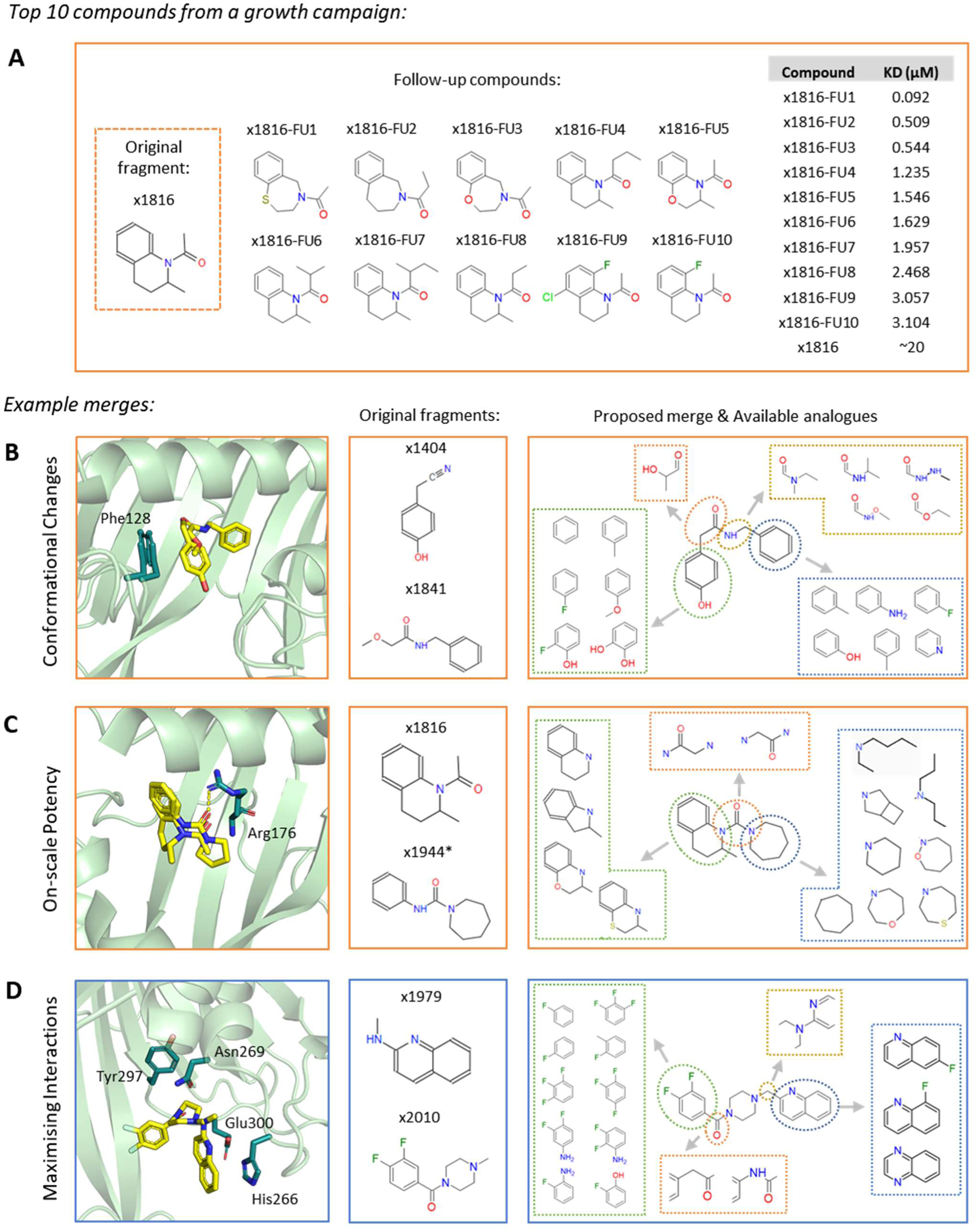
Opportunities for follow-up compound design: a growth campaign and example merges. **A.)** Top 10 compounds with the best affinity from a growth campaign aroundfragmentx1816.Chemical structures and KO from SPR data are shown. Examples of proposed fragment merges based on **B.)** conformational changes, **C.)** detectable inhibitory activity and **D.)** key interactions. Binding pose and structure of the original fragments, chemical structure of the proposed merge and closest purchasable compounds are shown.

Merging opportunities have also emerged which can explore the hypotheses outlined above. For example, to recapitulate a closed loop conformation, two fragments can be merged which both have caused the closed conformation of Phe128 on the 122-132 loop (conformation B), such as x1404 and x1841 (Fig. 8(B)). Alternatively, to explore the large shifts within the N-terminal domain, fragments x1437 and x1290 can be merged combinatorially.

When looking at compounds with measurable inhibition activity, most have bound at the distal end of the herbicide pocket and fall into one of two aforementioned pharmacophores (those that hydrogen bond to Arg176 and resemble current herbicides, and bicyclic compounds resembling 1,8-naphthyridine). Compounds of each group are quite similar but the combination of the two yields some promising structures such as x1944 and x1816 (Fig. 8(C)).

Lastly, when trying to maximise the interactions made by the follow-up compound, we picked sets of overlapping fragments that interact with key residues of the site and together span the largest area. For example, the joining of the deep and shallow sites within the active site through a merge between x1979 and x2010 could maintain hydrogen bonds with Asn269, salt bridge to Glu300 and crucially, π-stack with the catalytic His266 as well as expand to the hydrophobic network of Tyr297, Ile140 and Val142 (Fig. 8(D)). Inside the herbicide site expansion of the distal end will lead to recapitulation of the interactions made by current herbicides while the proximal end contains newly identified key interactors such as Gln111, Met146, Tyr297 and Asn269.

## Discussion

FatA’s conformational flexibility may be how it has adapted to receive large substrates and, in turn, may be exploited by inhibitors. The N-terminal half of FatA which is responsible for substrate binding^20^ is much more dynamic than the catalytic C-terminal half as evident from the large shifts induced by some ligands, the dynamic nature of the nearby loop 122-132 and the extensive side chain movement. What remains to be seen is if binding of the substrate forces these movements in an induced-fit model or whether FatA samples various conformations which are then selected by its substrate or its ligands. Inhibitors bound at the herbicide site may not only sterically exclude the substrate but also lock FatA in a closed conformation so that the fatty acid cannot enter the substrate access channel at all. These observations can be implemented to guide follow-up compound design through prioritisation of fragments, which caused more radical movements and a combination of fragments, which have caused the similar structuring of the mobile elements to promote the resulting compound to do the same.

The herbicide binding site, named herein for its binding to all known herbicides, is both highly populated and effective. The site is located deep within the central cavity formed between the beta-sheet and the alpha-helix of the N-terminal side of FatA. It has evolved to receive fatty acids and is predictably highly hydrophobic. Despite this, fragments traversed the long channel to make mostly hydrophobic interactions, although H-bonds with Arg176 and Tyr217 were also highly observed. Notably, it is fragments which bind here that cause the large conformational changes and structuring of the 124-129 residues. Additionally, the majority of fragments with on-scale potency against FatA bind at this site.

The fragment screen has revealed orthogonal opportunities for novel inhibitor development. Some of our fragments bind at the herbicide binding site but explore novel interactions and space when compared to current herbicides. Their further development would alleviate mutational pressure from the residues currently key to herbicidal interactions. We have also found two other sites which show promise, including one fragment (x1168/x1231) which binds at both and has shown detectable activity. The active binding site is closer to the surface and thus more accessible than the herbicide site, and it is likely that bigger ligands will be required to produce a greater inhibitory effect. However, we have found fragments that interact with the active site residues themselves and thus show great promise as starting points. The dimer site is also interesting since dimerization has been shown to be necessary for FATs’ activity^22^. Perhaps merged compounds can pry the two chains apart or affect catalysis allosterically, although there is a danger of stabilising the dimer instead. Since this site is located mainly in the C-terminal half of one chain, and that half is much less dynamic, the entropic cost of binding is likely to be less here.

The large number of bound fragments has readily lent itself to growing and merging opportunities, even when inspected by eye. Merging especially has been shown to be an effective way of quickly accessing on-scale kinetics and potency^34–37^. SAR exploration may then be achieved by purchasing the merged compounds and their analogues. Moreover, a small elaboration campaign based on the growth of an active x1816 fragment has already yielded an improvement in affinity from ∼20μM to ∼80nM. It may also be beneficial to test prospective leads against FatB since inhibition of both enzymes is necessary for lethality^2,18,19^.

In summary, conducting a crystallographic fragment screen enabled us to find many new starting points for future inhibitor development. Concurrently, obtaining more than a hundred crystal structures of FatA revealed conformational flexibility of its substrate-binding N-terminal half. This shined a light on possible mechanisms of substrate recognition and inhibition. These insights can be applied during rational follow-up compound design, thus demonstrating the utility of structural biology as a tool in herbicide discovery and development.

## Methods

### Protein Expression and Purification

pET-24a(+) plasmid containing codon optimised *Arabidopsis thaliana* FatA sequence was supplied by Syngenta. The N-terminal chloroplast transit peptide (residues 1-74) had been removed and a C-terminal His6 tag was added. See Supplementary Table 1 for the gene and protein sequence.

To express FatA, the plasmid was transformed into BL21(DE3) *Escherichia coli* and grown overnight at 37°C on lysogeny broth (LB) agar containing kanamycin (50 μg/mL). Several transformed colonies were inoculated into super optimal broth with catabolite repression (SOC) media containing kanamycin (50µg/mL) and grown overnight at 37°C. 10mL of starter culture was used to inoculate 1 litre of terrific broth containing 10mL of glycerol and 50µg/mL kanamycin and incubated at 37°C, 200 rpm until an optical density of 1. Temperature was reduced to 18°C and after 1 hour, 0.5mM isopropyl β-d-1-thiogalactopyranoside (IPTG) was used to induce protein expression. Cells were grown overnight at 18°C, 200rpm before being harvested by centrifugation and frozen at -80°C.

The cell pellet was resuspended in 25mM HEPES pH 7.5, 500mM NaCl, 0.5mM tris(2-carboxyethyl)phosphine (TCEP), 25mM Imidazole and 0.03 μg/mL benzonase. Cells were disrupted by sonication and the lysate was clarified by centrifugation at 30,000 × *g*. Supernatant was loaded onto a column packed with Ni Sepharose 6 Fast Flow resin (Cytiva, 17531802) equilibrated in 25mM HEPES pH 7.5, 500mM NaCl, 0.5mM TCEP, 25mM Imidazole, 10% glycerol. After washing, the protein was eluted with 25mM HEPES pH 7.5, 500mM NaCl, 0.5mM TCEP, 500mM Imidazole, and 10% glycerol. The protein was concentrated using a 10kDa molecular weight cut-off centrifugal concentrator (Sartorius, VS2002), and applied to an SRT-10 SEC-300 column (Sepax, 225300-21230) equilibrated in 25mM HEPES pH 7.5, 150mM NaCl, 0.5mM TCEP. The protein was concentrated to 21 mg/mL, flash-frozen in liquid nitrogen and stored at -80°C.

### Crystallisation and Structure Determination

Crystals were obtained through the sitting drop vapour diffusion method using crystallisation conditions previously identified by Syngenta. Crystals were grown at 20°C in SWISSCI three-well plates (SWISSCI, 3W96T-UVP) with 30μL of reservoir solution containing 0.1M MES pH 6.85 and 1.6M ammonium sulphate. Crystallisation drops consisted of 200nL of protein (8.3 mg/mL), 100nL of reservoir solution and 20nL of seeding stock. Crystals could be harvested after one day and did not require additional cryoprotectants. Seeding stock was used to improve the consistency of nucleation. To make the seeding stock, several crystals were crushed using a melted tip of a glass Pasteur pipette, transferred into 50μL of reservoir solution, crushed by vortexing around 10 times for 10 seconds with a seed bead (Fisher Scientific, 12398637), incubating briefly on ice in between, and used at 1:10 dilution.

Data was collected on beamlines I03 and I04 at Diamond Light Source and processed using Diamond’s fully automated processing pipelines which include Xia2, DIALS, autoPROC and STARANISO with the default settings^38–41^. Further processing was done in CCP4i2^42^. The dataset was phased with a FatA structure provided by Syngenta using Phaser^43^ and refined with Refmac5^44^. Model building was done in Coot^45^. The structures of FatA in the *apo* state and bound to an unidentified ligand have been deposited in the PDB under the codes 9HRR and 9HRQ respectively.

### Modelling of FatA, ACP and 18:1 Fatty Acid

FatA crystal structure from x1747 was chosen since it has the most open 122–132 loop. Since the loop in chain A was unstructured, chain B was copied into its place to make a symmetric molecule. Position of ACP was predicted in complex with an *apo* FatA dimer using ColabFold and then superimposed onto x1747 in PyMOL^46,47^. The 2FatA:2ACP model was then minimised using FastRelax in PyRosetta^48^. Position of the substrate was modelled based on 4KEH, a cross-linked structure between fatty synthase dehydratase, FabA, ACP and sulfonyl-3-alkynyl crosslinking probe (1R3)^32^. 4KEH and 2FatA:2ACP model were aligned in PyMOL. The sulfanyl 1R3 ligand was modified to be two linked moieties, phosphopantetheine (PNS) and a second moiety, either a transition state compound (custom structure) or oleoic acid (OLA). The structure was then minimised again. The model has been deposited in GitHub (https://github.com/matteoferla/A-thaliana-FatA-modelling) and Zenodo (https://doi.org/10.5281/zenodo.14413325).

### Crystallographic Fragment Screening, Data Processing and Analysis

Two fragment libraries were screened at XChem^26^, Diamond Light Source: Enamine’s essential fragment library (320 fragments) and Diamond-SGC-iNEXT Poised (DSipoised) library including the EUbOpen extension (781 fragments). *Apo* crystals were obtained as described above. Two crystallisation conditions were tested in a 100-fragment pre-screen: 0.1M MES pH 6.85, 1.6M ammonium sulphate and 0.1M sodium cacodylate pH 6.85, 1.6M ammonium sulphate. All fragments were in DMSO at 500mM. 15nL of fragment stocks were transferred into the 300nL crystallisation drops using an ECHO liquid handler (Labcyte), equating to a final concentration of 23.8mM and DMSO concentration of 5%. After incubating at 20°C for 2 hours the crystals were mounted using the Crystal Shifter (Oxford Lab Technology) and flash cooled in liquid nitrogen.

Data was collected on beamline I04-1 at Diamond Light Source and processed using Diamond’s automated processing pipelines as described above. Further processing was done in XChemExplorer which generates electron density maps with DIMPLE, generates ligand restraints using AceDRG or grade, creates an *apo* ground state model and uses PanDDA to find ligand-binding events within the bound state models^49–52^. Ligand hits were built using PanDDA-generated event maps, successive rounds of refinement with Refmac and model building with Coot^44,45^. Coordinates, structure factors, and PanDDA event maps were deposited in the PDB under group deposition ID G_1002328.

Molecular interactions were analysed using the Hit Interaction Profiling for Procurement Optimisation (HIPPO) Python package, Protein-Ligand Interaction Profiler (PLIP) web service, Schrödinger’s Maestro software suite and cross-validated using PyMOL^47,53–55^.

B-factor and RMSD analysis was performed using PyMOL and Knime^47,56^.

### FatA Enzymatic Assay

Compounds were tested for *in vitro* inhibition of Arabidopsis FatA using an assay for detection of the CoA product released following hydrolysis of oleoyl-coenzyme A (OCA) used as a surrogate substrate. Standards and test compounds were solubilised in DMSO and applied in 1ul to the plate well. The assay was carried out as follows: a reaction mixture consisting of 12nM FatA in assay buffer (50mM Tris HCl pH 8.0, 150mM NaCl, 12nM beta-casein (Sigma C6905)) was combined with test compounds in black 384-well microtitre plates (Greiner, Fluotrac 200, 781076) in a final reaction volume of 70µl. Plates were incubated for 15 min at room temperature with mixing (500 rpm). The plate was read (excitation 390nm, emission 470nm) using a Tecan Infinite plate reader to check for autofluorescent test compounds. The enzyme reaction was then initiated by the addition of OCA (Sigma O1012) to 10µM final assay concentration (added as 10µl of 80µM stock solution in assay buffer) and incubated for a further 30 min at room temperature with mixing at 500rpm. The reaction was terminated by addition of CPM solution (7-diethylamino-3-(4-maleimidophenyl)-4-methylcoumarin, Sigma (96669)) at 20µM final assay concentration (added as 10µl of 180µM stock solution in detection buffer (50mM Tris HCl pH 8.0, 150mM NaCl, 50% ethanol)). Following 10 min incubation in the dark at room temperature, the adduct generated by reaction of CPM with CoA was measured by recording the fluorescence intensity as above. Test compounds were assayed in an 8 rate dose response with a top rate of 25ppm using 1 in 5 dilutions. The standard, CSCA651647, was tested in an 8 rate dose response starting at 1ppm using 1 in 5 dilutions. Untreated, maximum, controls used 1ul of DMSO and minimum controls had a final assay concentration of 1ppm CSCA651647.

Data was analysed by nonlinear regression using GraphPad Prism (version 10.1.2 for Windows, GraphPad Software, Boston, Massachusetts USA, www.graphpad.com). The activity data was normalised using the fluorescence intensity from the standard (minimum control) and untreated (maximum control) samples. A curve was fitted using non-linear regression (log(inhibitor) vs. response - variable slope (four parameters)), which enabled extraction of the IC50 values.

### Surface plasmon resonance

FatA was immobilized on an NTA sensor chip (GE Healthcare) using a Biacore 8K, captured via His-tag and simultaneous amine coupling on EDC/NHS-activated carboxyl groups (Immobilization buffer: 25mM HEPES pH 7.4, 150mM NaCl, 1mM TCEP, 0.05% Tween 20). Equilibrium binding experiments were performed at 20°C using a Biacore 8K instrument (GE Healthcare) with 25mM HEPES pH 7.4, 150mM NaCl, 1mM TCEP, 0.05% Tween 20, 2% DMSO as running buffer for compound testing. Compounds were injected over the chip in 2-fold dilution series either in multi-cycle mode (60s/180s association/dissociation time) or single-cycle kinetic mode (160s/1800s association/dissociation time). Indicative KD values were obtained by steady-state affinity fitting to a 1:1 binding site model.

Compounds were drawn using ChemSketch^57^.

## Data Availability

The crystallographic coordinates and structure factors of FatA in the *apo* state and bound to an unidentified ligand have been deposited in the PDB under the codes 9HRR and 9HRQ respectively. The resulting structures from the XChem fragment screen have been deposited in the PDB under the group deposition ID G_1002328. Crystal structures of FatA bound to cinmethylin, oxaziclomefone and methiozolin have been deposited in the PDB with the codes 9GRR, 9GS1 and 9HMT respectively. The model of the biological assembly of FatA bound to 18:1 FA-ACP has been deposited in GitHub (https://github.com/matteoferla/A-thaliana-FatA-modelling) and Zenodo (https://doi.org/10.5281/zenodo.14413325). The authors declare that all other data supporting the findings of this study are available within the paper and its supplementary information files.

## Acknowledgements

The authors would like to thank Jo Mattocks in Protein Science at Syngenta for protein production and purification. We would also like to acknowledge the Diamond Light Source for beamtime on beamlines I03, I04 and I04-1 under proposals MX28172, MX34598, LB30602 and LB36049 as well as access to the fragment screening facility XChem and for usage of the EEF and DSi-Poised libraries. Finally, we would like to thank Dr. Dennis Wegener and his team at Evotec - Hamburg for SPR support.

## Author contributions

E.K. protein production, crystallography, XChem fragment screening, data analysis, wrote the first draft of the manuscript. F.v.D., N.P.M., M.G.M., K.S.E, L.O.V. supervised the research, read and edited the manuscript. J.C.A., X.N., C.W.E.T. read and edited the manuscript. M.G.M initial crystallography. N.P.M. and M.G.M. x1816 follow-up SAR exploration. M.F. protein production support. X.N., L.K. crystallographic support. X.N. PDB deposition support. M.P.F. created the substrate-bound model of the biological assembly. C.W.E.T., D.F., J.C.A. XChem support. M.P.F. and L.O.V. chemical proofreading. P.H.H. enzyme assay. M.W. data analysis of the interactions.

## Materials and correspondence

Correspondence and requests for materials should be addressed to F.v.D.

