## Supplementary Information for "Crystal structure of Fatty Acid Thioesterase A bound by 129 fragments provides diverse development opportunities"

**Supplementary Table 1**. Gene and protein sequence of FatA used.

**Gene Sequence**

atgggtagcctgaccgaggatggcctgagctacaaagagaagttcgtggtgcgcagctacgaagtgggcagtaataaaaccgccaccgtggagaccatcgcaaatctgctgcaggaagttggctgcaaccatgcacagagcgtgggttttagcaccgacggtttcgccacaacaacaaccatgcgcaagctgcatctgatctgggtgaccgcccgcatgcatatcgagatctacaagtacccggcctggggtgacgtggttgaaatcgaaacctggtgccagagcgaaagtcgtattggtacccgccgtgactggattctgaaggatagcgtgaccggcgaagttaccggccgtgccaccagcaagtgggtgatgatgaaccaggatacccgccgcctgcagaaagtgagcgatgacgtgcgcgatgagtatctggtgttttgcccgcaagagccgcgcctggcatttccggaggagaacaatcgcagcctgaaaaagatcccgaagctggaagacccggcccagtatagtatgattggcctgaaaccgcgccgcgcagatctggatatgaatcagcatgttaataatgtgacctatattggctgggtgctggaaagtatcccgcaggagattgtggacacccacgaactgcaggttatcaccctggactatcgccgtgaatgccagcaggacgatgtggtggatagcctgacaaccaccaccagcgaaattggtggcaccaatggcagcgcaaccagcggcacccagggtcataatgacagccagttcctgcatctgctgcgtctgagcggcgatggccaggaaattaatcgcggcaccaccctgtggcgcaaaaaaccgagtagccatcatcatcaccaccactaa

**Protein Sequence**

MGSLTEDGLSYKEKFVVRSYEVGSNKTATVETIANLLQEVGCNHAQSVGFSTDGFATTTTMRKLHLIWVTARMHIEIYKYPAWGDVVEIETWCQSEGRIGTRRDWILKDSVTGEVTGRATSKWVMMNQDTRRLQKVSDDVRDEYLVFCPQEPRLAFPEENNRSLKKIPKLEDPAQYSMIGLKPRRADLDMNQHVNNVTYIGWVLESIPQEIVDTHELQVITLDYRRECQQDDVVDSLTTTTSEIGGTNGSATSGTQGHNDSQFLHLLRLSGDGQEINRGTTLWRKKPSSHHHHHH

**Supplementary Note 1.** Analysis of fragment screening data.

A pre-screen using 100 fragments from the Enamine Essential Fragment (EEF) Library was conducted on the 0.1M MES pH 6.85 and 1.6M ammonium sulphate condition and the 0.1M sodium cacodylate pH 6.85, 1.6M ammonium sulphate condition. Since both resulted in unique hits, both have been included in the “Fragment Screen Full” sheet in the Supplementary Information.xlsx. A Soaking Fail (where the crystal deterioration prevented mounting after soaking with the fragment) is denoted by “fail” in the “Mounted?” column. A Diffraction Fail (where the crystal did not diffract after mounting) is denoted by a “fail” in the “Diffraction?” column. For the first 100 compounds, a failure is only classified as such if it is present in both pre-screens (denoted as “double fail”). The rest of the EEF and the DSI-poised libraries were screened as single repeats.

**Supplementary Note 2.** Analysis of SPR data.

60 analogues of the fragment x1816 were screened using SPR by Evotec. All compounds were analysed by stead state affinity fit. Some compounds could then be further analysed by kinetic fit. Compounds x1816-FU1, x1816-FU2, x1816-FU3, x1816-FU4, x1816-FU39 and x1816-FU55 were repeated again in duplicate. Therefore, when comparing all the analogues, the final KD used and presented in Figure 8(A) is either the average of the repeats where a repeat was performed, the kinetic KD where a kinetic fit was performed or if neither is present, the affinity KD.

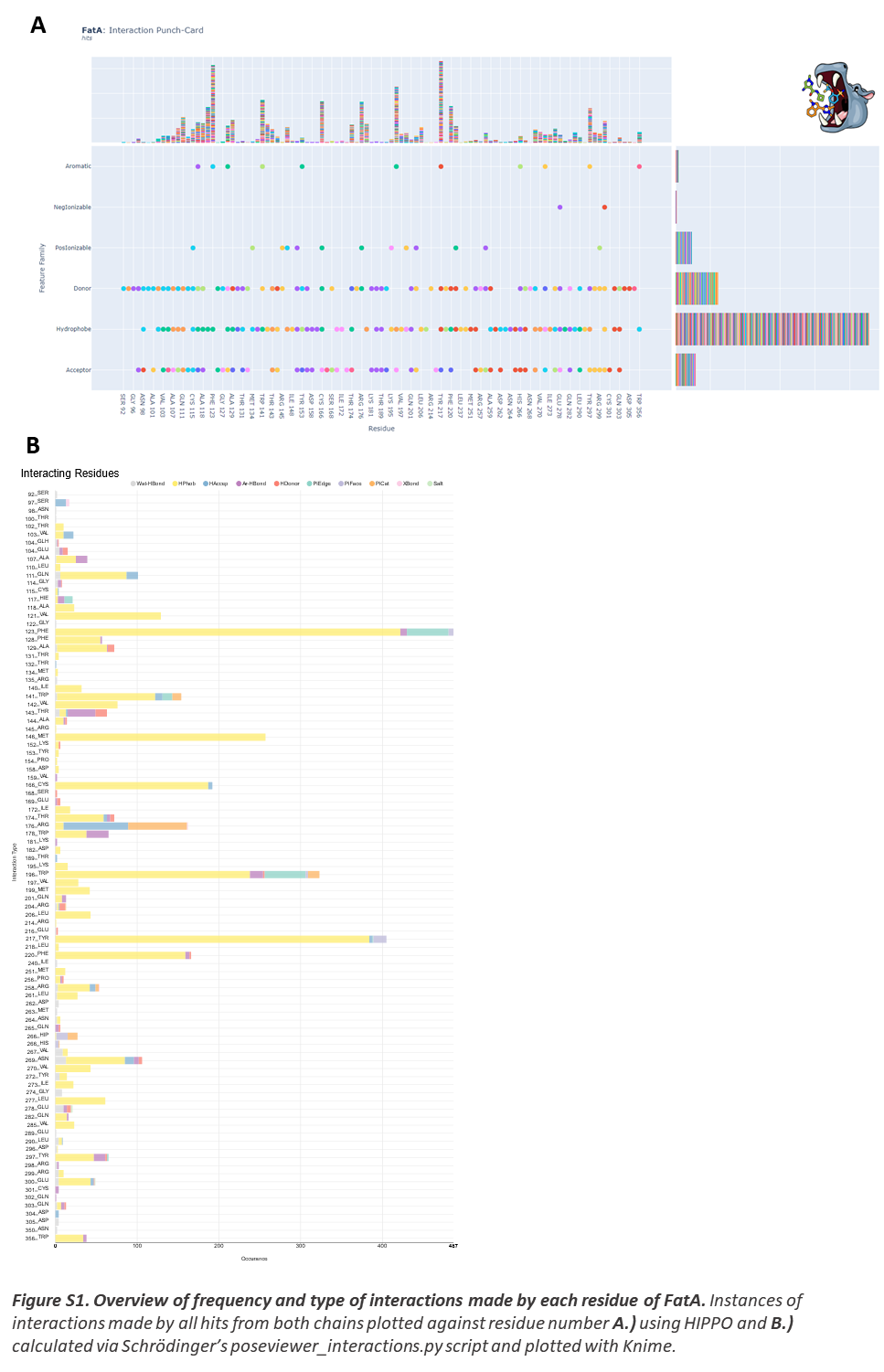

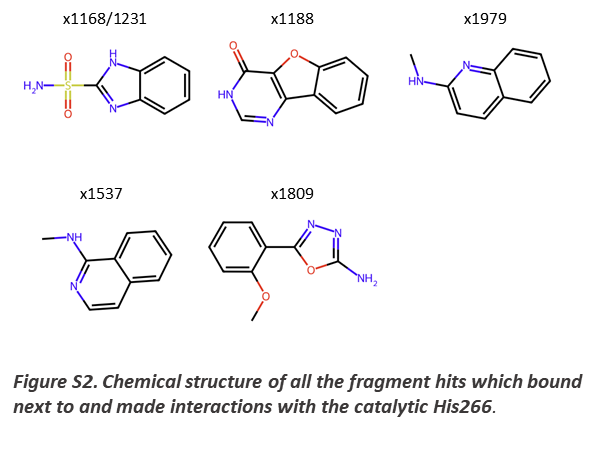

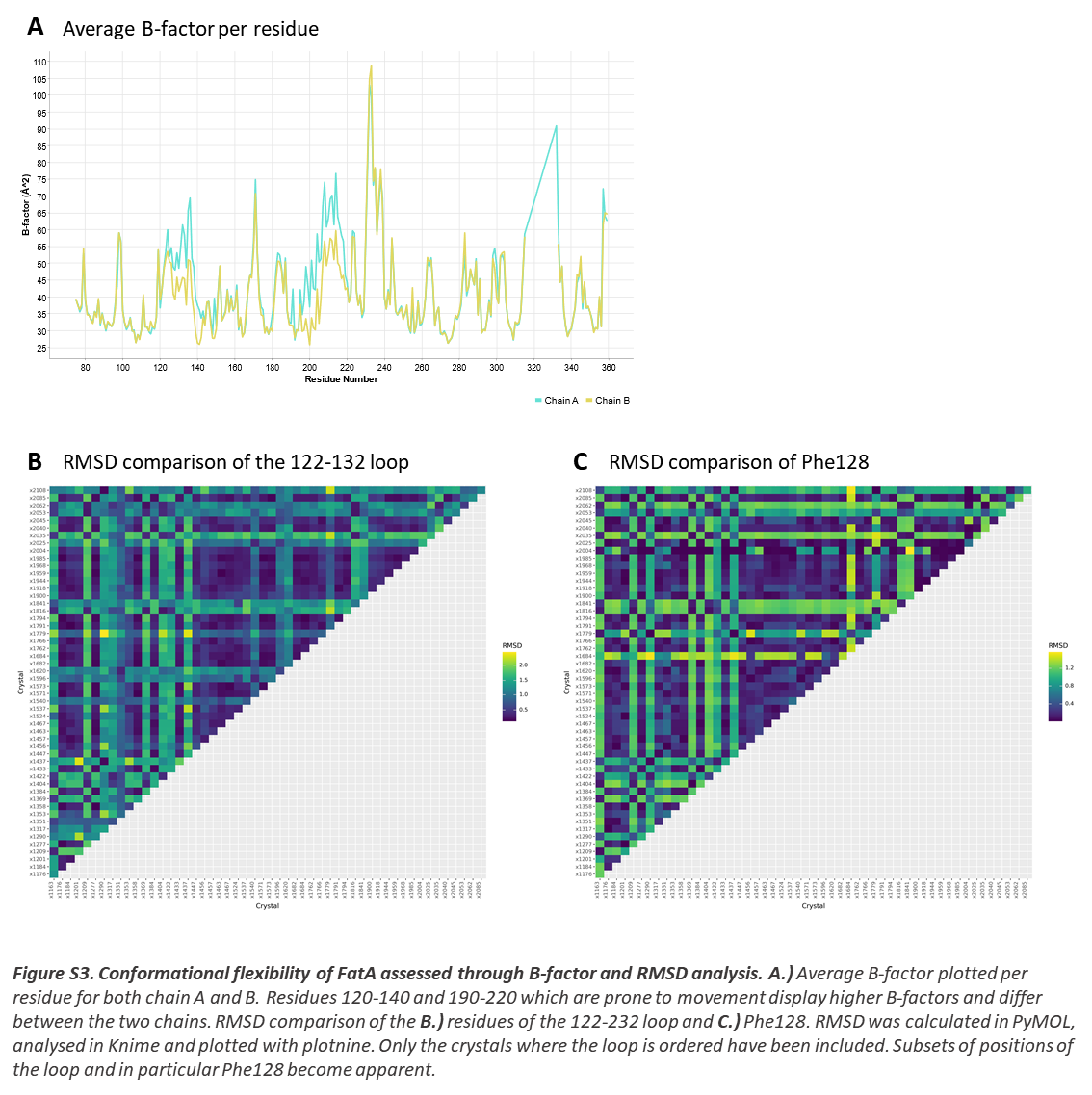

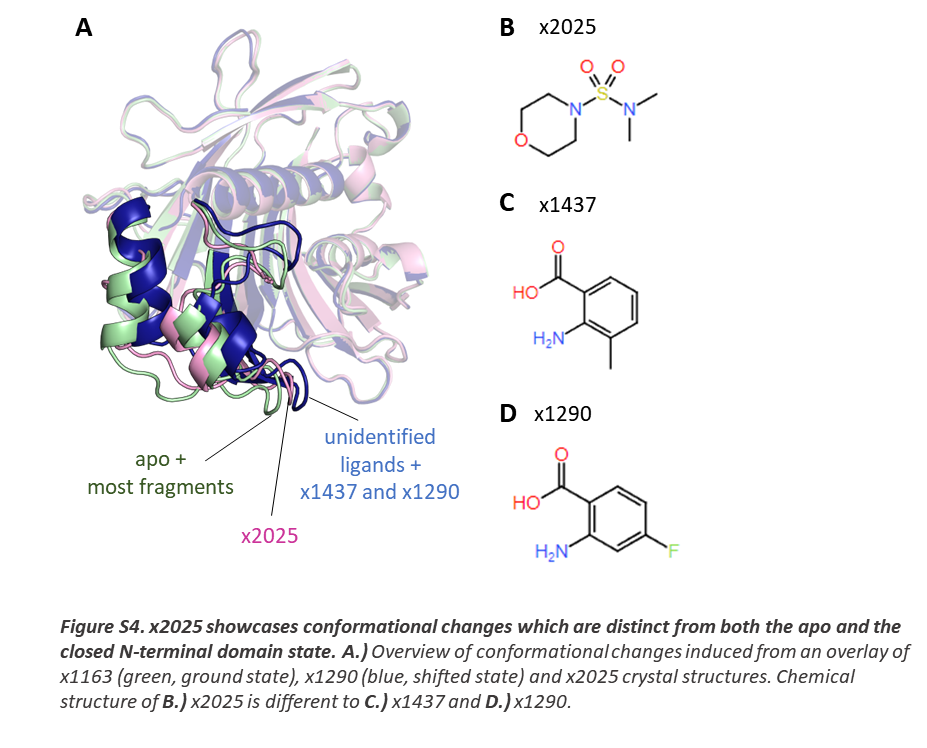

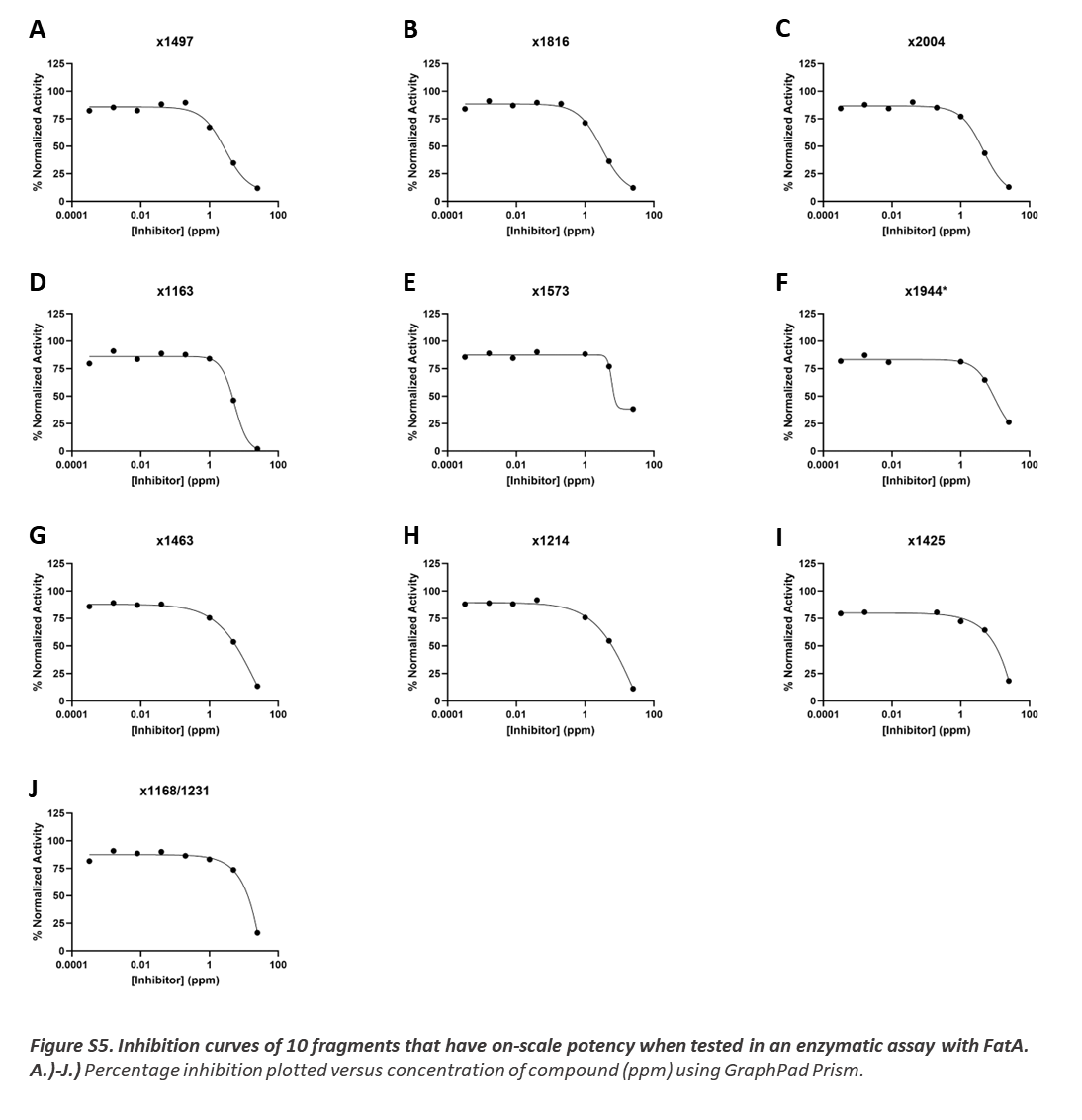
